# An innate immune activation state prior to vaccination predicts responsiveness to multiple vaccines

**DOI:** 10.1101/2021.09.26.461847

**Authors:** S. Fourati, LE. Tomalin, MP. Mulè, DG. Chawla, B. Gerritsen, D. Rychkov, E. Henrich, HER. Miller, T. Hagan, J. Diray-Arce, P. Dunn, The Human Immunology Project Consortium (HIPC), O. Levy, R. Gottardo, M. Sarwal, JS. Tsang, M. Suárez-Fariñas, B. Pulendran, RP. Sékaly, SH. Kleinstein

## Abstract

Many factors determine whether an individual responding to vaccination will generate an immune response that can lead to protection. Several studies have shown that the pre-vaccination immune state is associated with the antibody response to vaccines. However, the generalizability and mechanisms that underlie this association remain poorly defined. Here, we sought to identify a common pre-vaccination signature and mechanisms that could predict the immune response across a wide variety of vaccines. We leveraged the “Immune Signatures Data Resource” created by the NIH Human Immunology Project Consortium (HIPC) to integrate data from 28 studies involving 13 different vaccines and associate the blood transcriptional status of 820 healthy young adults with their responses. An unsupervised analysis of blood transcriptional profiles across studies revealed three distinct pre-vaccination states, characterized by the differential expression of genes associated with a pro-inflammatory response, cell proliferation, and metabolism alterations downstream of NFκB and IRF7. Innate and adaptive immune cell subset-specific genes were also associated with the three pre-vaccination states. Importantly, individuals whose pre-vaccination state was enriched in pro-inflammatory response genes known to be downstream of NFκB tended to have higher serum antibody responses one month after vaccination. A supervised analysis of the same data resulted in a single classifier, also enriched for NFκB regulated genes, that predicted the antibody response across most of the vaccines. Projection into single-cell RNA-sequencing data suggested that this pre-vaccination state was attributable to the signature of activation of non-classical monocytes and myeloid dendritic cells. Transcriptional signatures of acute responses to bacterial and not viral infections were enriched in the high pro-inflammatory pre-vaccination state and also included NFκB regulated genes. The pro-inflammatory pre-vaccination state was highly reminiscent of the innate activation state triggered by TLR ligands or adjuvants. These results demonstrate that wide variations in the transcriptional state of the immune system in humans can be a key determinant of responsiveness to vaccination. They also define a transcriptional signature NFκB activation at baseline, that is associated with a greater magnitude of antibody response to multiple vaccines, and suggest that modulation of the innate immune system by next-generation adjuvants targeting NFκB before vaccine administration may improve vaccine responsiveness.

## Introduction

Prophylactic vaccination is a cost-effective strategy to prevent or reduce the effect of viral and bacterial infections. Vaccine efficacy often varies in the population and can depend on by age^1^, sex^2^, ethnicity^3^ and genetics^4^. Human immune responses are also shaped by the environment, including previous pathogenic perturbation of the immune system. Indeed, pre-vaccination predictors of antibody response to specific vaccines such as influenza, yellow fever and hepatitis B vaccines have been identified^5–8^, including baseline predictive signatures of both influenza and yellow fever vaccines^9^. However, whether pre-vaccination markers exist for all vaccine platforms or if universal pre-vaccination markers of vaccine response can be identified have not been addressed for a large number of vaccines.

To define the biological signatures associated with the induction of protective immune responses induced by vaccination, high-throughput transcriptomic technologies (microarray and RNA sequencing) have been used to profile the peripheral blood cells of vaccine recipients. Paired with the use of machine-learning techniques, previous studies have identified signatures (*i*.*e*. sets of genes) of vaccine conferred protection and/or of protective antibody responses to immunization. For example, different aspects of pre-vaccination states, including the frequency of B cell subsets as well as the expression of genes related to B cell receptor signaling and antigen processing predicted antibody response to influenza, yellow fever and hepatitis B vaccinations^5,8–10^. In contrast, pre-vaccination expression of genes related to granulocytes and interferon (IFN)-stimulated genes have been associated with a poor response to hepatitis B vaccination^5,11^. Genes related to proliferation and inflammatory responses were also shown to be expressed at a higher level by participants with a poorer response to the influenza vaccine^6,12^ and the malaria vaccine^13^. However, a common pre-vaccination signature shared by all of these vaccines has yet to be identified. Moreover, some of the biological pathways identified showed opposite associations with response between vaccines (*e*.*g*., IFN signaling is negative predictor of antibody response for hepatitis B^11^ but type I IFN genes are positive predictor of antibody response for influenza and yellow fever vaccination^9^), or between studies for the same vaccine (*e*.*g*., B cell signaling for influenza vaccination^10,12^). The interpretation of these differences can often be complicated by not only the vaccine type, but also factors such as geographic region (e.g., whether the targeted pathogen is endemic vs. not), age, and different genes in the same pathway (or geneset) driving the association signals. The interaction of those various factors is complex and they could thus confound smaller-size cohort studies. Meta-analyses, leveraging information from many cohorts, have the advantage of an increased power to detect pre-vaccination signatures predictive of antibody responses to vaccines while minimizing the effects of co-founding variables (e.g., age, ethnicity, geographical region).

Identifying a universal pre-vaccination signature predictive of antibody responses to vaccines and understanding the biological pathways associated, and therefore potentially required for, inducing a protective humoral response following vaccination in healthy adults may lead to more effective strategies (*e*.*g*., administration of immunomodulators) to enhance vaccine response^14^. Those new strategies may particularly benefit the most vulnerable populations, including infants, the elderly, and immunosuppressed individuals.

Here, we show that a common pre-vaccination peripheral blood transcriptional signature is predictive of antibody responses across 13 different vaccines. Functional annotation of this signature shows enrichment of effector genes of pro-inflammatory responses and pre-exposure sensing of ligands associated with bacterial infections. Single-cell transcriptomic data showed that non-classical monocytes and myeloid dendritic cells as the likely source of this pre-vaccination signature. The overlap between this predictive signature and the transcriptomic signature following Toll-like receptor (TLR) stimulation or adjuvant treatment suggests that a state of natural adjuvantation is associated with better responses to vaccination.

## Results

### Heterogeneity of transcriptional profiles pre-vaccination

Transcriptomic profiles of whole blood and peripheral blood mononuclear cells of 820 healthy adults aged 18 to 55 before and after vaccination were collected from publicly available databases (refer to as the “Immune Signatures Data Resource”^15^). Several vaccine platforms ranging from live-attenuated viruses (*i*.*e*. yellow fever, smallpox and influenza vaccines), inactivated viruses (*i*.*e*. influenza vaccine) and glycoconjugate vaccines (*i*.*e*. pneumococcal and meningococcal vaccines) were included in this dataset (**Figure 1A-B**). We assessed the contribution of different socio-demographic (age, biological sex, ethnicity) and experimental (vaccine platform, time after vaccination) variables on the variance in the transcriptomic data (**Figure 1C**). Age (14%), timepoints (9%) and vaccine (9%) explained only a small fraction of the variance observed in the transcriptomic data; over 62% of the variance between samples could not be explained by any of the recorded clinical and experimental variables. To understand the source of the variance between participants, we restricted our analysis to the pre-vaccination timepoints (**Figure S1**). We used hierarchical clustering to identify subgroups of participants with similar transcriptomic profiles pre-vaccination.

**Figure 1.**
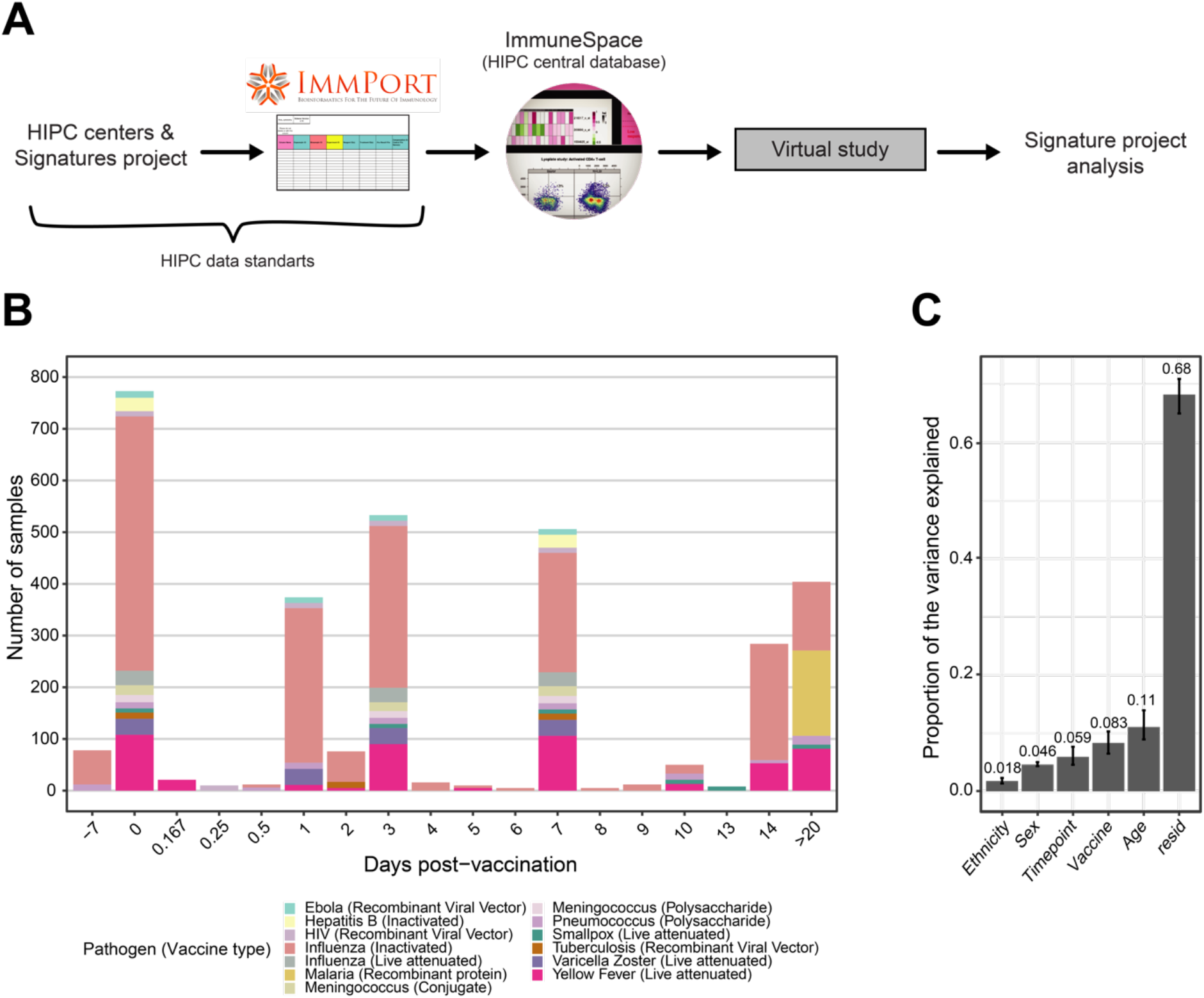
Creation of a combined dataset of transcriptional responses to vaccination across diverse pathogens and vaccine types **(A)** Flowchart describing the collection, curation, standardization and preprocessing steps leading to the creation of the vaccine transcriptomics compendium. **(B)** Histogram of the timepoints pre- (days -7 and 0) and post-vaccination (days > 0) available in the immune signature data resource. In the plot, each vaccine is represented by a different color, while the size of the bar is proportional to the number of samples with available transcriptomic data. Only healthy adults, aged 18 to 50 years old, with available pre-vaccination data were included in the resource. **(C)** Principal variance component analysis was used to estimate the proportion of the variance observed in the transcriptomic data that can be attributed to clinical (age, sex, ethnicity) and experimental variables (time after vaccination, vaccine). The proportion of the variance that could not be explained by those variables is depicted by the residuals (resid). Confidence intervals (95%, percentile method) and bar height (mean) were computed from 4000 bootstrap replicates.

### Pre-vaccination states of the immune system modulate the transcriptional response to vaccines

Hierarchical clustering (an unsupervised method) followed by identification of the number of clusters by the Gap statistic identified three groups of participants (i.e. states) based on their pre-vaccination expression of genesets included in the MSigDB hallmark genesets ^16^ and blood transcriptomic modules ^17^ (**Figure 2** and **Figure S2A**). Neither age, sex, nor pre-existing antibody levels to the immunogen were associated with the differences in gene expression observed in these three states (**Figure S2B**). Using samples collected 7 days before vaccination and those just before vaccination (Day 0) from the same participants (n=74), we confirmed the stability over time of these transcriptomic profiles (**Figure S2C**).

**Figure 2.**
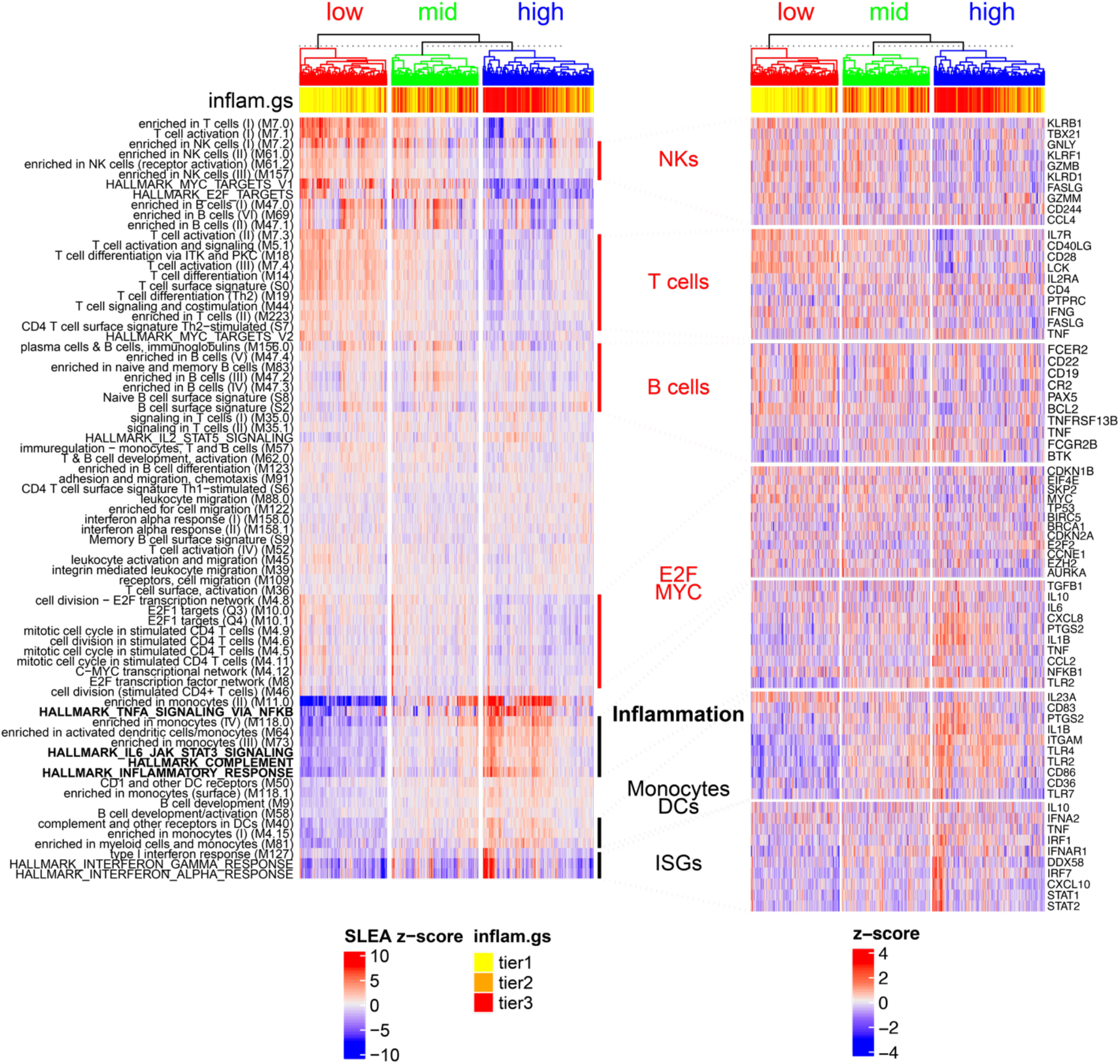
Participants have distinct pre-vaccination transcriptomic profiles. Hierarchical clustering (Euclidean distance metric and complete linkage agglomeration method) of pre-vaccination samples (day -7 and day 0) based on the expression of the blood transcriptomic modules (BTM) and hallmark genesets. The overall transcriptomic activity of genesets/modules were estimated using the SLEA method^32^. Three groups of participants/states can be identified by cutting the dendrogram. Average SLEA score of the four hallmark inflammatory genesets (bold row labels; inflam.gs), discretized in tertiles, is shown as sample annotation. One state with heightened pre-vaccination inflammatory pathways (i.e. high), one with low levels of inflammatory pathways (i.e. low) and one intermediary state (i.e. mid). Expression of genes part of the hallmark and BTM genesets are presented in a heatmap (right side).

One state showed heightened expression of transcriptomic markers of monocytes and dendritic cells, IFN-stimulated genes (ISGs) and pro-inflammatory genes and thus was designated a high inflammatory (inflam.hi) state. Transcriptomic markers of monocytes and dendritic cells induced in the inflam.hi state included several innate immune sensors (TLR1, TLR2, TLR4) and concomitantly genes of the TLR4 signaling cascade (TLR4, LY96, DNM3, PLCG2) (**Figure S2D**). The type I IFN signaling cascade was also an important feature of the inflam.hi state. Receptors upstream of the IFN pathways (IFNA2, IFNAR1, IFNAR2, TYK2), nucleic acid sensors that trigger this pathway (DDX58, TRIM25, MAVS, TRAF6, TANK), and transcription factors that regulate the expression of ISGs (STAT1, STAT2, IRF1, IRF7) were all upregulated in the inflam.hi state compared to the other two states. The NFκB pathway, a hallmark of inflammation, and its target genes, including pro-inflammatory cytokines (TNF, IL6, IL1B) and their receptors (TNFRSF1A) or effector molecules regulated by NFκB, including the metalloprotease ADAM17 that cleaves the ectodomain of TNF-*α*, were all induced in the inflam.hi stat. Likewise, the IL-6 signaling pathway (IL6R, JAK2, STAT3), a pathway that triggers the proliferation of activated B cells, was induced in the inflam.hi state. Moreover, several genes of the inflammasome complex and IL-1 signaling, also downstream of NFκB, were also upregulated in this state of subjects, including IL1A, IL1B, IL1R1 and IL1RAP. Altogether, this state was characterized by genes and pathways involved in pro-inflammatory processes common to nucleic acid-sensing, which could promote the development of an immune response to vaccines.

A second state showed lower expression of the above-listed pro-inflammatory genes and pathways (i.e. NFκB and ISGs) when compared to the first state (**Table S1**). This state was designated as the low inflammatory (inflam.lo) state. Heightened expression of transcriptomic markers of natural killer cells, T cells, B cells and target genes of the transcription factors E2F and MYC both involved in the upregulation of cell proliferation and cell metabolism were features specific to the inflam.lo state. Transcriptomic markers of natural killer (NK) cells induced in the inflam.lo state included cell surface markers of NK cells (KLRD1, KLRB1), effector molecules of cytotoxic function (GZMB, FASLG, CASP3), and genes of the IL12 signaling cascade (IL12RB1, STAT4). Transcriptomic markers of T cells expressed in the inflam.lo state included members of the IL2 signaling cascade (IL2RA, IL2RB, LCK), CD28 dependent PI3K/AKT signaling cascade (CD28, CD80, PIK3CA, PIK3R1, PIK3R3, AKT3) and IL7 signaling cascade (IL7, IL7R); the latter two pathways being involved in the maintenance of the naïve T cell pool. Transcriptomic markers expressed by B cells of the inflam.lo state included cell surface receptors (CD79A, CD79B, CD22, CD19) and kinases (FYN, BTK) of the BCR signaling complex. Known target genes of E2F and MYC induced in the inflam.lo state include cell cycle and proliferation regulators (MYC, CDKN2A, AURKA) and cell metabolism (LDHA, MTHFD2, TYMS). Altogether, this state was characterized by the lack of expression of genes downstream of innate sensing (i.e. IFNs and NFκB target genes), while their transcriptomic profiles showed that cells of the adaptive immune system were activated and engaged in an ongoing immune response ^18^.

Finally, a third state showed a mixed transcriptomic profile between low and inflam.hi states and was designated as the mid inflammatory (inflam.mid) state. T cells, NK cells and B cell-specific genes were upregulated in these participants compared to the inflam.hi state and higher levels of pro-inflammatory genes are found in this state compared to the inflam.lo state (**Table S1**).

### Immune cell frequencies vary between the pre-vaccination states

Flow cytometry (n=164) and immune cell deconvolution ^19,20^ were used to determine if the three pre-vaccination inflammatory states were driven the frequency of different innate and adaptive immune cell subsets (**Figure S2E**). The inflam.lo state showed an increased frequency of naive B cells (CD19+CD27-IgG-IgA-cells with heightened expression of ABCB4, ADAM28 and BACH2), which is in line with the above-described gene expression profiles. CD8+ T cells (CD3+CD8+CD45RA+ cells with heightened expression of CRTAM, PIK3IP1, TRAV12-2) were also more prevalent in this state. In contrast, the inflam.hi states showed an increase in Monocyte frequencies (19% of immune cells in inflam.hi versus 16% in inflam.lo), in line with the results from the transcriptomic profiling (**Figure 2**). To assess whether the change in gene expression between the three states could be explained solely by the difference in immune cell frequency, differential expression analysis was performed, adjusting for the immune cell frequency, and re-identified inflammatory genes as markers of three states (**Table S2**). This analysis suggests that the difference in inflammatory gene expression between the three states could not be explained by differences in cell frequencies alone and confirmed the differential transcriptomic activity of those inflammatory genes between states.

### The pre-vaccination states are associated with the early gene expression response to vaccines

Next, we evaluated the impact of the pre-vaccination inflammatory states on the magnitude and kinetics of post-vaccination transcriptional responses. The pre-vaccination inflammatory states explained 12.5% of the variance in gene expression observed pre- and post-vaccination (**Figure S3A**). Participants from the inflam.hi state showed reduced vaccine-induced expression of pro-inflammatory pathways (*e*.*g*., complement pathway, IL6 signaling pathway) at Days 1 and 3 post-vaccination when compared to the participants from the inflam.low (log_2_ fold-change (log_2_FC) < −1.46; Wilcoxon-rank sum test: *p*<0.0106) and inflam.mid (log_2_FC < −0.643; Wilcoxon rank-sum test: *p*<0.0996; **Figure 3A** and **Figure S3B**) states. By day 7, levels of the pro-inflammatory pathways returned to pre-vaccination levels in all three states. Similarly, participants from the inflam.hi state showed reduced expression of ISGs at day 1 post-vaccination when compared to the inflam.low (log_2_FC=−2.81; Wilcoxon rank-sum test: *p*=8.08×10^−4^) and inflam.mid (log_2_FC=−1.54; Wilcoxon rank-sum test: *p*=0.0996; **Figure 3B** and **Figure S3C**) states. The inflam.hi state participants also had a dampened B cell signature on day 7 and beyond compared to the inflam.low state (log_2_FC=−0.866; Wilcoxon rank-sum test: *p*=1.87×10^−4^; **Figure 3C** and **Figure S3D**). The levels of B cell markers returned to pre-vaccination levels by day 7 in the inflam.lo group contrary to the inflam.hi where B cell markers were sustainably induced compared to pre-vaccination levels (**Figure S3E**). Similarly, T helper 2 cell markers, necessary to mount an humoral response, were induced at day 7 post-vaccination in the inflam.hi group but not in the inflam.lo (**Figure S3F**). The inflammatory states affected the magnitude of the transcriptomic changes triggered by the vaccines, specifically at the earliest time points. However, we did not observe kinetic differences (i.e. delays in gene expression) between the three states.

**Figure 3.**
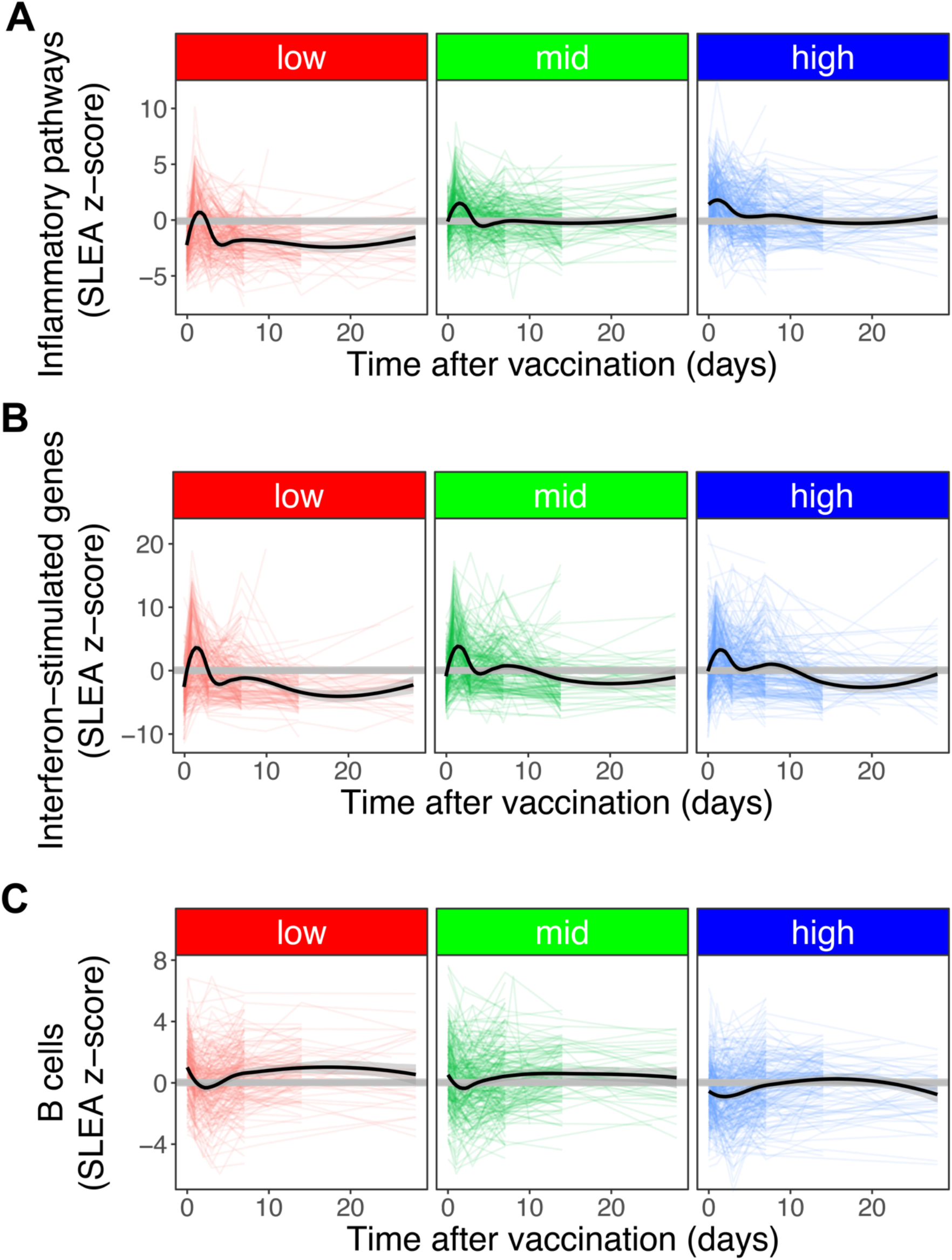
Kinetic of response dictated by pre-vaccination profiles. Line plot showing the expression of inflammatory pathways (**A**), ISGs (**B**) and B cells (**C**) as a function time, separately for participants with low, mid or high pre-vaccination inflammation. Each colored line corresponds to one participant, LOESS regression was used to determine the average expression per pre-vaccination state (black lines).

### Identification of universal predictive signatures of antibody responses to vaccination

We then assessed the association between the pre-vaccination states and antibody responses triggered by all 13 vaccines and measured 1 month (Day 28) after immunization. Participants from the inflam.hi state showed significantly higher antibody responses across all vaccines compared to participants of the inflam.lo state (log_2_FC=1.58, Wilcoxon rank-sum test: *p*=0.0161, **Figure 4A**). The association between the inflammatory states and antibody response was stronger for influenza inactivated vaccines but remained significant for the remaining vaccines (**Figure S4A**). The inflammatory states tended to be associated with antibody response measured beyond Day 28 but did not reach significance (**Figure S4B**). Taken together, there is an association between pre-vaccination immunological states and vaccine-induced antibody response.

**Figure 4.**
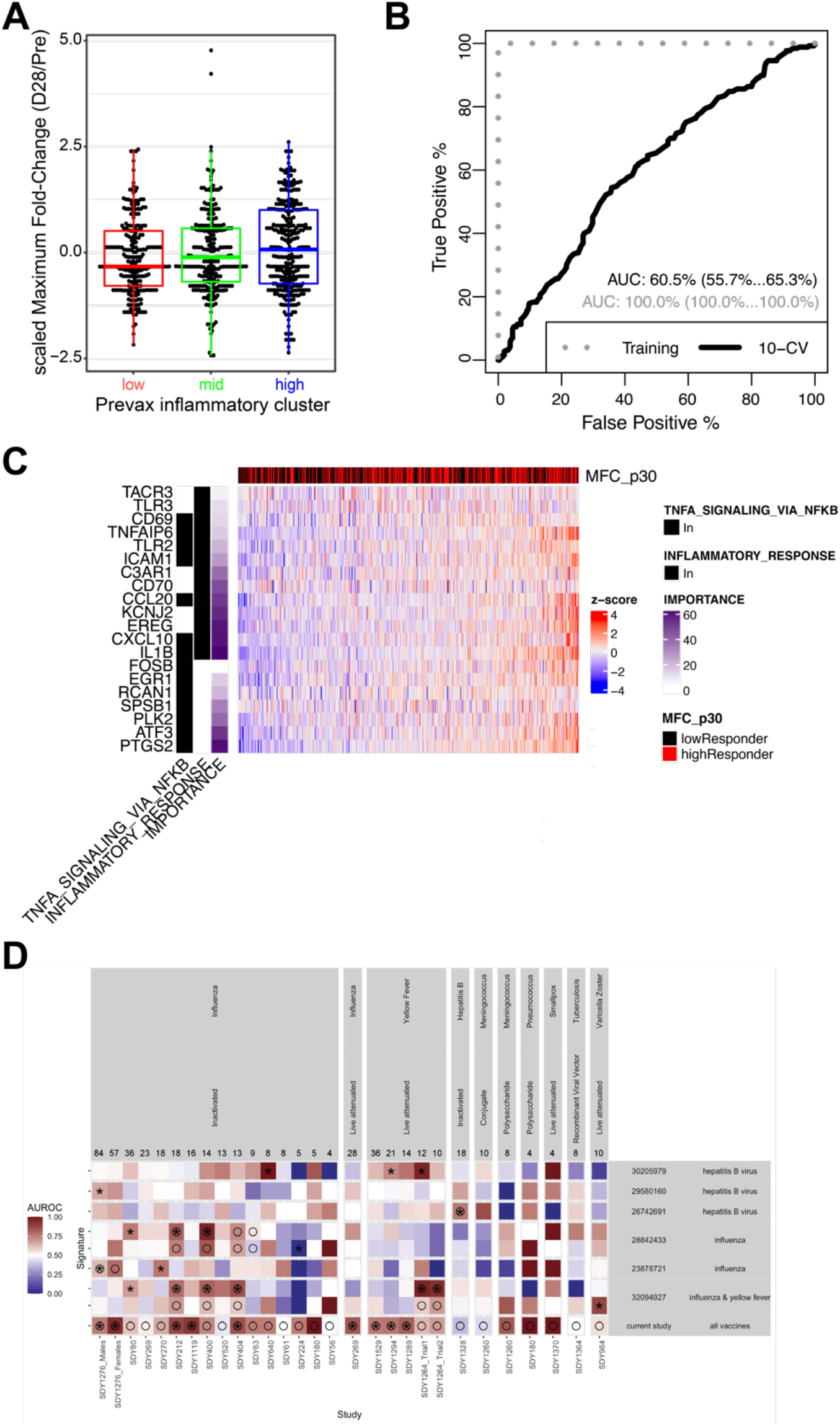
Prediction of antibody response by pre-vaccination profiles. (**A**) Boxplot of the maximum fold-change (MFC) between antibody responses at day 28 over pre-vax as a function of the pre-vaccination inflammation states. The MFC was scaled to a mean of 0 and a standard-deviation of 1 across vaccines (sMFC). A Kruskal-Wallis test was used to assess differences in antibody response between states and resulted in a p-value of 0.0323. (**B**) A supervised approach consisting in building an random-forest classifier was adopted to distinguish high vaccine responders (3^rd^ tercile) from low vaccine responders (1^st^ tercile) at day 28. The accuracy of the ensemble model was assessed by 10-fold cross-validation (10-fold CV). The receiver operating characteristic (ROC) curve is presented along with the area under the ROC curve (AUC) with 95% CI estimated from the 10-fold CV. (**C**) Top predictive genes/features included in the ensemble overlap inflammatory genes identified in the unsupervised approach (Fisher’s exact test: p < 3.93e-4). Heatmap showing the expression of the overlapping genes pre-vaccination. Samples columns are ordered by increasing levels of expression of the inflammatory genes. A Wilcoxon-rank sum test was used to assess association between the inflammatory signatures and high/low antibody response and resulted in a p-value of 4.25e-6. (**D**) Comparison of the gene signature in this work compared to previously identified pre-vaccination signatures of vaccine response. MetaIntegrator was used to apply each previously published pre-vaccination signatures of vaccine response, as well as the signature identified in this work, to the transcriptomic studies of “Immune Signatures Data Resource”. Circles correspond to studies used to train those pre-vaccination signature while asterisks indicate better than random identification of high-responder on a transcriptomic study.

To complement the unsupervised approach, we used a supervised approach to identify genes that are predictive of high (top 30%) versus low (bottom 30%) antibody response to vaccination. We trained a random forest classifier that predicts vaccine-specific antibody responses based on pre-vaccination gene-expression profiles. This classifier achieved an area under the ROC curve of 60% as estimated by 10-fold cross-validation (**Figure 4B**). The accuracy of the classifier was significant for the vaccines with the greatest number of samples (Influenza inactivated: n=476; *p*<2.95×10^−33^; Yellow fever: n=96; *p*=1.32×10^−3^) and deteriorated for vaccines with smaller sample sizes (**Figure S4B**, n<30; *p*>0.322). We did not observe any significant association between misclassification and the sex, age, ethnicities or geographical locations of the participants, suggesting that the classifier accuracy is not affected by those parameters. For example, the yellow fever vaccine recipients included in the immune signature dataset originated from five cohorts recruited in the United States, Canada, Switzerland, Uganda and China. The supervised classifier was significantly associated with high vaccine response in all cohorts except the one from the United States. The immune signature datasets also include vaccines that were administered intramuscularly, intravenously or intranasally (*e*.*g*., FluMIST), and the inflammatory signatures were predictive independently of the route of vaccination.

The top 200 predictive genes selected by their importance in the classifier were enriched for inflammatory markers (Fisher’s exact test: p=4.25×10^−6^; **Figure 4C**). Inflammation markers (identified in **Figure 2**) that contributed to the classifier predictions included several pro-inflammatory cytokines and chemokines (CCL20/MIP3a, CXCL10/IP-10, IL1B), receptors involved in innate immune signaling (TLR2, TLR3, CD70/TNFSF7), mediators of complement activation (C3AR1, ICAM1) and pro-apoptotic effector molecules (CASP7, CASP10). The classifier was compared to six previously identified pre-vaccination signatures of vaccine responses^5,6,9,11,21,22^. There was no significant overlap in gene content between the supervised classifier and the six previously identified pre-vaccination gene signatures (**Figure S4C**). Notably, the classifier developed here was the only one to predict antibody response across the majority of the vaccines tested. In contrast, most of the previously identified signatures, including the pro-inflammatory signature we previously identified that predicted influenza vaccination response^6^, were largely predictive for the vaccine types they have been trained on less on the remaining vaccine types (**Figure 4D**). Altogether, the signature identified here provides evidence that a specific inflammation signature pre-vaccination helps to mount a good antibody response across multiple vaccines.

### Etiology of the pre-vaccination inflammatory states

To identify the cells that potentially express the inflammatory genes included in the classifier of vaccine-induced antibody responses, we utilized CITE-seq data from PBMCs collected from 20 healthy participants prior to vaccination with an inactivated influenza vaccine ^9^. We tested if the inflammatory genes were elevated in a specific cell subset or if their expression reflected a heightened global state of immune cell activation pre-vaccination common to all subsets. We analyzed the expression of the inflammatory genes of the classifier of vaccine-induced antibody responses within clusters of single cells defined by the expression of 65 specific cell surface proteins **(Figure 5A** and **Figure S5)**. The inflammatory genes identified by the unsupervised and supervised analysis were highly enriched within the innate immune cell subsets compared to other cell populations, in particular within CD14^+^ CD16^-^ classical monocytes, CD14^-^ CD16^+^ non-classical monocytes, CD1c^+^ CD11c^+^ mDCs, and CD123^+^ CD303^+^ pDCs **(Figure 5B)**. These results suggested that the cellular source of the pre-vaccination activated state found through orthogonal supervised and unsupervised analysis was derived from innate immune myeloid cells (as suggested from **Figure 2**).

**Figure 5.**
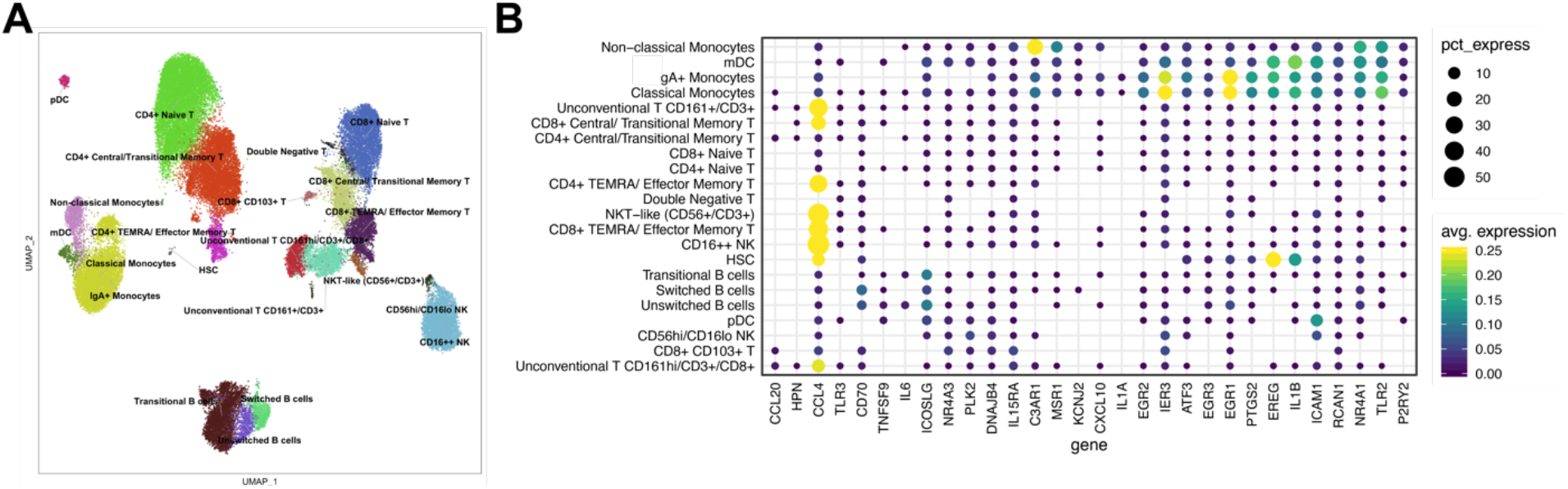
Pre-vaccination states in scRNAseq (**A**) UMAP of PBMCs from 10 high and 10 low responders profiled by CITE-seq^9^; subsets were identified based on surface protein expression (average dsb normalized protein expression within each cluster). (**B**) Single cell CITE-seq deconvolution of inflammatory genes identified by the unsupervised and supervised approaches as being associated with antibody response in the blood cells subsets. The color represents average log normalized expression within each cluster with scales clipped at a maximum of 0.25, and the dot size represents the percent of cells within that cluster with non-zero expression of the gene.

Pre-vaccination inflammation in seemingly healthy participants can result from a non-infectious etiology or from bacterial- or viral-induced proinflammatory responses. To identify the possible upstream signals associated with the inflammation described above, we used the 7-gene classifier described in Sweeney et al.^23^ to discriminate between inflammation caused by bacterial (classifier score above 0) or viral infections (classifier score below 0). Applying this classifier to our cohort of vaccinees showed that participants within the inflam-high state and the highest Ab response expressed genes associated with exposure to bacterial infections (**Figure 6A**).

**Figure 6.**
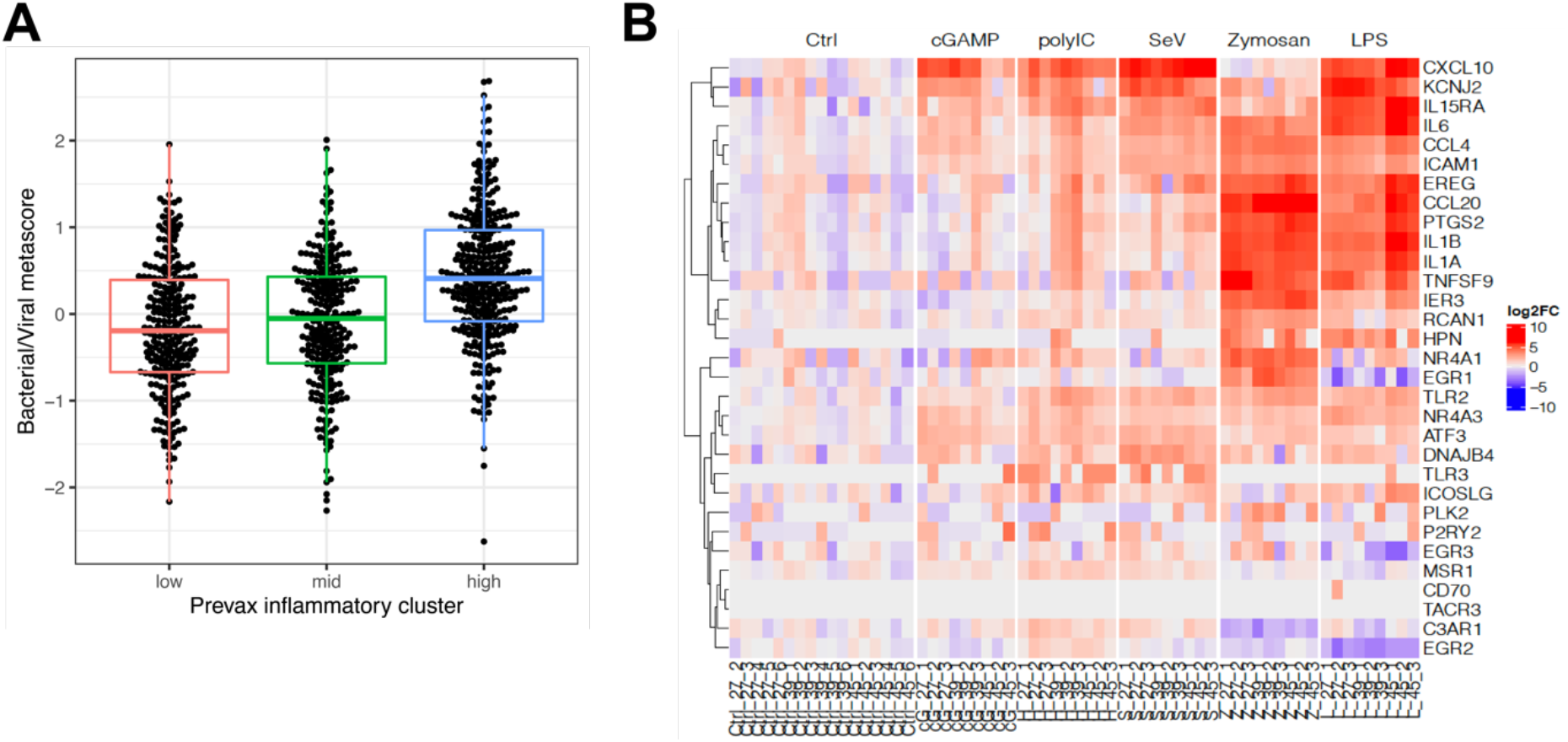
**(A)** Boxplot showing the Bacterial/Viral metascore^23^ as a function of the pre-vaccination inflammatory states. A Kruskal-Wallis test was used to assess the difference in bacterial/viral metascore between states and resulted in a p-value of 3.76×10^21^. **(B)** Gene expression of dendritic cells from three independents donors stimulated for 6 hours with five PRR ligands.

We further observed that one of the bacterial markers in this 7-gene classifier, TNIP1, is a known NFκB target and that the classifier score was positively correlated with an induction of NFκB target genes. This contrasts with IFI27, a ISG used as a viral marker in the 7-gene classifier, and that IFN targets negatively correlated with the bacterial/viral classifier score. Interestingly, vaccines that were correctly predicted by the classifier show a stronger induction of NFκB targets in high-responders than low-responders (**Figure S6A**; ex: Influenza inactivated: log_2_FC=2.48; Yellow fever: log_2_FC=0.743; Hepatitis B: log_2_FC=1.12). ISGs, downstream of IRF7, were also associated with a robust humoral response to most of the vaccines except vaccines using poxvirus vectors such as the Smallpox or Yellow fever vaccines; for which strong expression of ISGs were associated with hyporesponses (**Figure S6A**).

To confirm those results, we queried publicly available transcriptomic datasets related to bacterial inflammation^24^, viral inflammation^24^, PRR activation^25^ and antibiotic treatment^26^. Again, counter-intuitively, our inflammatory signature generated on healthy participants showed significantly overlapping with gene signatures from participants infected by *S. aureus* and *S*. *pneumoniae* compared to healthy participants and to peripheral mononuclear cells stimulated *in vitro* with the TLR2/6 ligand PAM2 (**Figure S6B**). Gene expression of dendritic cells stimulated with pattern recognition ligands (several of them used as vaccine adjuvants)^27^ showed strong induction of the inflammatory genes that were part of our classifier, suggesting that the heightened expression of those genes is a hallmark of a naturally adjuvanted immune system (**Figure 6B**).

## Discussion

In this manuscript, we characterize the inter-individual heterogeneity in the inflammatory state of the peripheral immune system pre-vaccination associated with vaccine response. We show that this heterogeneity is characterized by different transcriptional signatures, which are also associated with a distinct distribution of cell subsets pre-vaccination. Our results show that this heterogeneity is associated broadly with the relative magnitude of the antibody response to 13 different vaccines. Our work highlights the impact of the role of the pre-vaccination immune system and pre-sensitization of the innate immune system to pathogen-associated molecular patterns in priming the B cell response to vaccination. The universality of previous pre-vaccination signatures across vaccines and populations has not been established. The results presented here extend these earlier observations to more diverse vaccines and populations; more importantly, they provide a mechanistic framework that can lead to the selection of adjuvants most efficient at stimulating vaccine-induced protective immune responses.

The inflammatory signature identified in this work predicts antibody response with a significant accuracy across the 13 vaccines tested. Compared to previously identified pre-vaccination signatures of vaccine response, ours was thus the closest to a universal signature of vaccine response. This suggests that a set of genes and pathways associated with protective antibody responses following immunization are shared among vaccines. This is noteworthy as our analysis included a broad range of vaccines that engage several innate immune system cells and molecules. For example, the live attenuated yellow fever vaccine will engage TLR2 and TLR8 on mDCs, TLR7 and TLR9 in pDCs, and RIG-I/MDA5 ^28^, Smallpox virus will engage STING, whereas inactivated influenza vaccine will engage TLR7/TLR8.

The approach undertaken herein that consisted of training on all 13 vaccines distinguishes this work from previously published reports. This strategy is most likely the main factor contributing to the identification of this pan-vaccine classifier. Training this classifier on one vaccine type did not confer predictive power on distinct vaccine types irrespective of whether this was a live attenuated, inactivated, or subunit vaccine (data not shown). In contrast, the global classifier of vaccine responses identified herein performed as well as a classifier trained on a given vaccine and tested on that same vaccine. Our classifier performed better on vaccines with a greater sample size suggesting that the accuracy of our classifier could be improved by performing transcriptomic analysis of future studies for those vaccines where we were able to obtain a limited set of samples.

Our results show that qualitative and quantitative features, including transcriptional programs (MYC and E2F versus IFNs and NFκB target genes), can identify a pre-vaccination environment that will lead to heightened antibody response to vaccines. Expression of NFκB, the prototypic transcriptomic factor that controls the development of inflammatory responses, and its target genes are induced in the inflam.high state. NFκB is essential for driving the transcription of cytokines (*e*.*g*., TNF) and chemokines (*e*.*g*., CXCL10) that trigger cells of the innate and adaptive immune responses to migrate to sites of vaccination and differentiate into effector cells. Consistent with our previous reports on positive pre-vaccination signatures of antibody responses to vaccination^14^, upregulation of ISGs is a feature of this state of participants, including the master transcriptomic factor of the type I/type II IFNs cascades IRF-7. Type I and type II IFNs regulate genes involved in antigen processing and presentation. In contrast, inflam.lo participants demonstrated upregulation of transcriptional networks that highlight genes and pathways of T and B cell activation, proliferation while these same participants showed low NFκB and IRF7 expression.

These two pathways seem to be driven by acute response to bacteria (NFκB) or to virus (interferons) infections. Both signatures show synergy (additivity) with vaccines that trigger MyD88 and IRFs, suggesting that the activation of these pathways in innate immune cells will lead to more efficient priming of innate immune responses. Indeed, both TNF and the inflammasome are potent inducers of adaptive immune responses and are triggered by several adjuvants including Alum and MF59. Of note, presence of the IFN signature prior to vaccination can also be negatively associated with the antibody responses in live attenuated viral vaccines in some populations (yellow fever, Smallpox, dengue vaccine^29^). This inhibitory effect of IFNs may be due to their antiviral activity, which could limit viral replication and antigen production by vaccines.

The heightened transcriptional signature of inflammation-related genes pre-vaccination, confirmed to be stable over a week-long period could result from (i) host genetics; (ii) the environment, which includes diet, prior infection etc, and (iii) the microbiome. To the latter point, our previous work showed that TLR5-mediated sensing of flagellin in the gut microbiota promoted influenza vaccine specific antibody response by stimulating lymph node macrophages to produce plasma cell growth factors ^30^.

The inflammatory response has been linked to aging; a process that has been termed inflammaging. Compared to young adults, increased inflammation in the elderly has been reported to be associated with hyporesponse to vaccines. The inflammatory signature identified here was not associated with the humoral response to influenza, hepatitis B and varicella zoster vaccines in the elderly, suggesting that age-associated inflammation^5^ is different (i.e. lacking intersecting genes) from the inflammatory signals associated with vaccination-response in adults (18 to 55 years). This suggests that different types of inflammation can lead to different responses to vaccination. Indeed, we provide direct evidence that inflammation is heterogeneous across individuals and associated with vaccine responses. Importantly, we show that this inflammatory signature that was associated with response to vaccination overlapped with the inflammation triggered by exposure to bacterial byproducts including or by translocation of bacteria from the gut; the latter signature is different from the inflammation caused by non-infectious diseases, viral infection or antibiotic therapy. The inflammatory signature described in this work contains several TLR genes and appears to be concentrated in mDCs and non-classical monocytes. These leukocytes are different from those mediating other types of inflammation listed above, suggesting that the inflammation associated with vaccine response may result from different biological drivers than other types of inflammation. The prevalence of different types of inflammation is also suggested by the lack of overlapping genes with inflammation signatures in the elderly (inflammaging) which have been associated with poor vaccine response. In addition to a distinct relationship of inflammatory states to vaccine responses, additional factors contribute to age-specific immune responses to adjuvants and vaccines, including distinct PRR function with age^31^.

Our data show that higher frequencies of monocytes are observed in participants with high inflammatory responses (14% to 26% of immune cells in blood). In contrast, participants of the low inflammatory state demonstrated high frequencies of naive B cells and CD8 T cells. Although we observed differences in immune subset frequencies between the pre-vaccination states, those frequencies could not solely explain the differences in gene-expression observed between the pre-vaccination states, highlighting that in addition to differences in cellular composition of blood, pre-vaccination states also reflect differential transcriptomic activities associated with the state of the immune system pre-vaccination.

Participants from the inflam.lo states showed several marks of a distinctly activated immune system prior to vaccination, including a heightened expression of E2F and MYC transcriptomic program and heightened frequency of CD8+ T cells. In addition, the inferred frequency of CD8+ T cells from the deconvolution analysis was negatively correlated with Day 28 antibody response suggesting that participants of the inflam.lo states may have an activated/committed immune system prior to vaccination.

Strategies that directly impact pre-vaccination inflammation or modulate the pre-vaccination commensal bacterial flora impact the immune response to vaccination^14,26^. In this study, we observed similarities between the pro-inflammatory signature associated with vaccine response and the pro-inflammatory signatures induced by bacterial infection. Bacterial infections activate pattern recognition receptor signaling cascades, which will trigger the activation of the NFκB transcription factor complex and the induction of pro-inflammatory transcriptomic programs. The overlap between the pro-inflammatory signatures associated with vaccine response and following bacterial signaling was not specific to one bacterial species but was shared by different bacteria such as *S. aureus* and *S. pneumoniae*. The signature overlapped with that of the activation of PRRs by bacterial immunogens such as TLR1, TLR2 and TLR4. Interestingly, the expression of TLR1, TLR2 and TLR4 were identified in the pro-inflammatory signature that was associated with enhanced responses to vaccines. Moreover, engagement of other adjuvants, such as polyIC enhances the expression of PRRs and the induction of the same pro-inflammatory genes as those associated with robust vaccine responses. Among the 13 vaccines part of the immune signature dataset, only the hepatitis B vaccine was adjuvanted with aluminum hydroxide. For the other vaccines that did not use an adjuvant, having a pro-inflammatory signature pre-vaccination originating from dendritic cells and non-classical monocytes appeared to reflect an activated innate immune state and was associated with an enhanced humoral response post-vaccination.

In conclusion, the inflammatory signature identified herein predicts humoral response across diverse vaccines and provides a mechanistic framework that can lead to the selection of adjuvants most efficient at stimulating vaccine-induced protective immune responses. Whether the heightened inflammatory response is also associated with humoral response to current or future vaccines platforms (mRNA, nanoparticle, adenoviruses) not included in the meta-analysis, including vaccines against SARS-CoV-2, remains to be determined.

## Material and Methods

### Gene-expression Preprocessing

They included 2,949 samples from published studies and 228 samples not included in previously published studies. All these samples were assembled into a single resource (referred to as the immune signature dataset).

An extensive description of the preprocessing of microarray and RNA-Sequencing (RNA-Seq) datasets included in the immune signature dataset can be found at ^15^. Briefly, raw probe intensity data for Affymetrix studies were background corrected and summarized using the RMA algorithm. For studies using the Illumina array platform, background correct raw probe intensities were used. For RNA-Seq studies, count data was voom-transformed to mimic gene array expression intensities distribution. Expression data within each study is quantile normalized and log-transformed separately for each study.

### Batch correction

An extensive description of the across studies normalization used to correct for batch effect can be found at ^15^. Briefly, a linear model was fit using the pre-vaccination normalized gene-expression as a dependent variable and platform, study and cell types as independent variables. The estimated effect of the platform, study and cell types was then subtracted to the entire gene expression (pre- and post-vaccination) to obtain the batch corrected gene expression used for the analysis presented in this article. Principal variance component analysis (PVCA) was used to assess the effect of other phenotypic variables on the batch corrected gene expression. All the phenotypic variables were coded as categorical variables before the PVCA analysis; this includes the age imputed coded as 10-years intervals and the timepoints before and after vaccination left-censored at 20 days and coded as days from vaccination.

### Clustering of the samples

For functional characterization of the genes, we made use of known genesets from two sources: Hallmark collection from MSigDB (version 7.2) ^16^ and the blood transcriptomic modules (BTM)^17^. Overall activity on each geneset/pathway was estimated for each sample using Sample-Level Enrichment Analysis (SLEA)^32^. Hierarchical clustering using Euclidean distance and complete linkage was used to cluster samples. The resulting dendrogram was cut to generate three clusters of samples. The three clusters were designated as low-, mid-, high-inflammatory clusters based on the average SLEA z-score of four hallmark inflammatory genesets (HALLMARK_INFLAMMATORY_RESPONSE, HALLMARK_COMPLEMENT, HALLMARK_IL6_JAK_STAT3_SIGNALING and HALLMARK_TNFA_SIGNALING_VIA_NFKB). Hallmark and BTM genesets were grouped based on their name and description into markers of seven cell subsets or canonical pathways (T cells, NK cells, B cells, Monocytes/DC, Inflammation, E2F/MYC and ISGs). Canonical genes of those seven cell subsets or canonical pathways were identified by looking at the genes part of the genesets annotated to those cell subsets or canonical pathways and ranking them based on the number of GeneRIF entries associating them to cell subsets or canonical pathways. The expression of the top 10 genes annotated to those seven cell subsets and canonical pathways are presented in the gene-level heatmap.

### Antibody response

Maximum fold-change was calculated for all participants with neutralizing antibody response, HAI, or IgG levels measured by ELISA ^15^.

### Identification of high and low responders

The maximum fold-change (MFC) between day 28 (+/-2 days) and pre-vaccination tiers was used to quantify the antibody response to vaccination. To minimize the difference in antibody response between studies (due to differents vaccines, different techniques used for antibody concentration assessment), the high and low responders were identified for each study separately by selecting the participants with MFC above the 70th percentile as high responders and participants with MFC below the 30th percentile as low responders.

### Strategy for identifying a gene signature predictive of vaccine response using pre-vaccination transcriptomic profiles

In order to evaluate if participant-specific transcriptomic profiles taken pre-vaccination were predictive of antibody response 28 days post-vaccine, we set out to develop predictive models using the random forest algorithm. The training set included participants achieving a high or low antibody response (n=522) based on the discretization of maximum fold-change (MFC_p30) and top 500 varying genes as input. The predictive model was trained to minimize the Brier score and tuning parameters were estimated based on 10-fold cross-validation. In this final model, the performance was assessed using 10-fold cross-validation using standard performance metrics including auROC, Accuracy, PPV, NPV, Sens, Spec, as well as Brier score.

MetaIntegrator was used to apply previously identified pre-vaccination signatures of vaccine response and the predictive model identify in this work to the different studies part of ‘Immune Signatures Data Resource’^33^. The area under the receiver operating curve (auROC) was used to assess the accuracy of the signatures.

### CITE-Seq analysis

CITE-seq data consisting of pre-vaccination PBMC samples from participants in SDY80 were downloaded from ^9^. Cell type annotations used in this analysis are the ‘high resolution’ annotations from Kotliarov et. al, these clusters were derived from graph-based clustering using Seurat^34^ directly on a Euclidean distance matrix of surface protein expression. CITE-seq surface protein data was normalized and denoised using dsb^35^. UMAP embeddings were calculated using the umap python package^36^. A Wilcoxon Rank Sum test with a minimum proportion of expressing cells of 0.2 and a log fold-change threshold of 0.3 was used to test genes from the high inflammatory state between different clusters using Seurat (all genes shown in the heatmap in Figure 5).

### Comparison with other pre-vaccination signatures

The bacterial/viral classifier was applied to the immune signature dataset by averaging the expression of the bacterial infection markers (HK3, TNIP1, GPAA1, and CTSB) and subtracting the average expression of the viral infection markers (IFI27, JUP, and LAX1); a resulting score above or equal 0 was considered more similar to bacterial infection while a score below 0 more similar to viral infection.

## Data availability

All data used in this study are available in ImmuneSpace (www.immunespace.org/is2.url).

## Code availability

R code used to generate the figures presented in the paper can be found at (www.immunespace.org/is2.url).

## Acknowledgments

This research was performed as a project of the Human Immunology Project Consortium (HIPC) and supported by the National Institute of Allergy and Infectious Diseases. This work was supported in part by NIH grants U19AI118608, U19AI128949, U19AI090023, U19AI118626, U19AI089992, U19AI128914, U19AI128910, U19AI118610, U19AI128913, and the Intramural Program of NIAID and NIH institutes supporting the Trans-NIH Center for Human Immunology (CHI).

