## Supplemental Figures for "An innate immune activation state prior to vaccination predicts responsiveness to multiple vaccines"

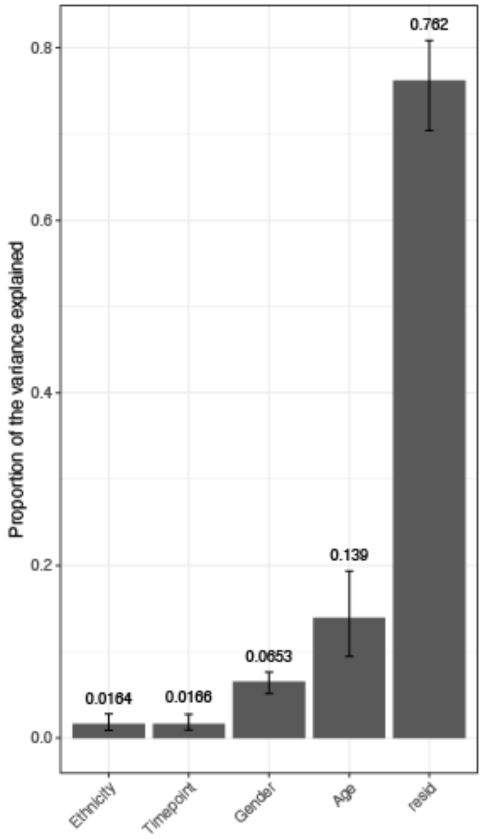


**Figure S1.** Principal variance component analysis using pre-vaccination transcriptomic expression. All phenotypic variables were coded as categorical variables. 95% intervals of confidence were calculated by bootstrapping the samples. resid: residuals.

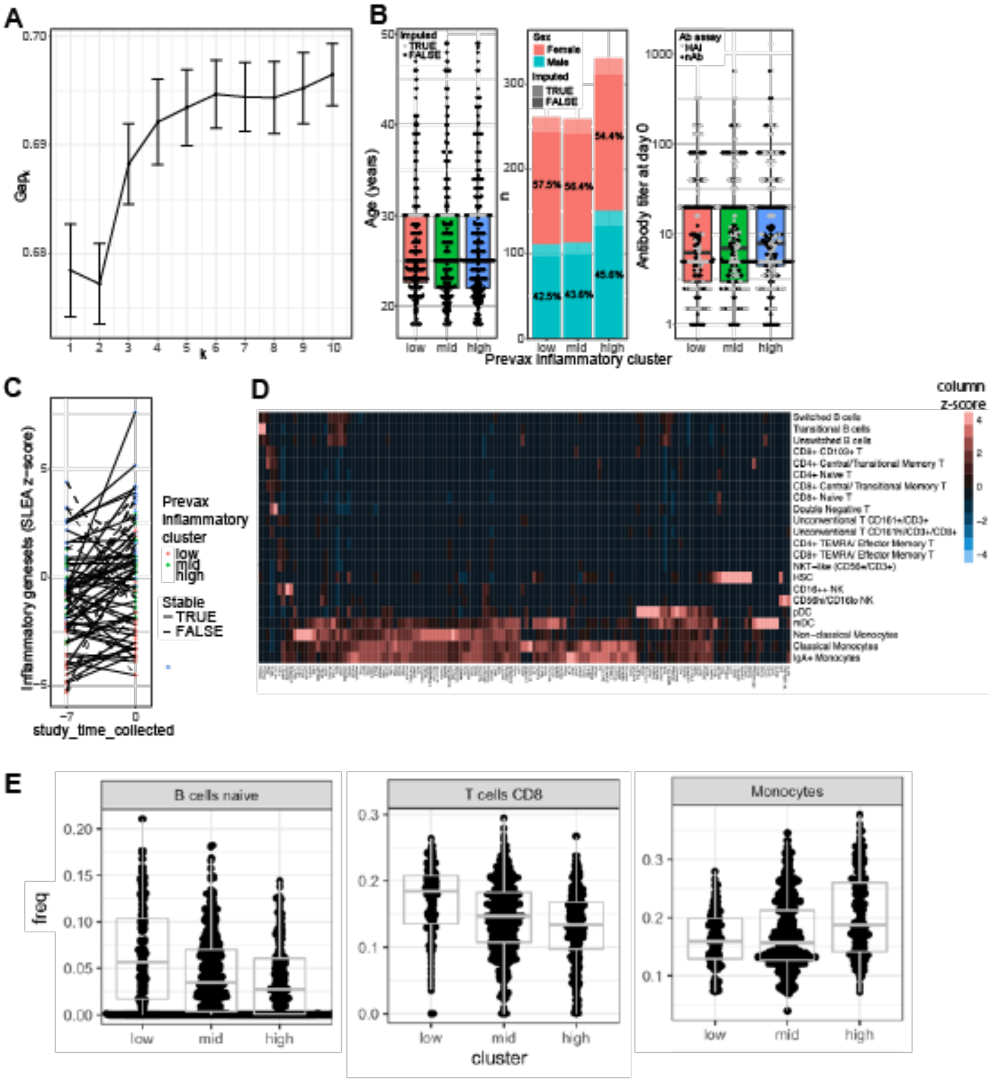


**Figure S2. (A)** Gap statistic. **(B)** Boxplot of the age as function of the inflammatory clusters. **(C)** Lineplot showing the differences in expression of inflammatory pathways between 7 days before vaccination compared to just before vaccination. **(D)** Monocytes/dendritic cell markers expression in blood of pre-vaccinated individuals. PBMCs from 10 high and 10 low responders profiled by CITE-seq[1]. The heatmap shows the average expression of monocyte and dendritic cell markers identified in bulk meta-analysis associated with the high inflammatory state, scale shown is the z score of the gene across protein based cell subsets. **(E)** Frequencies of immune cells, estimated by deconvolution, in the three pre-vaccination states

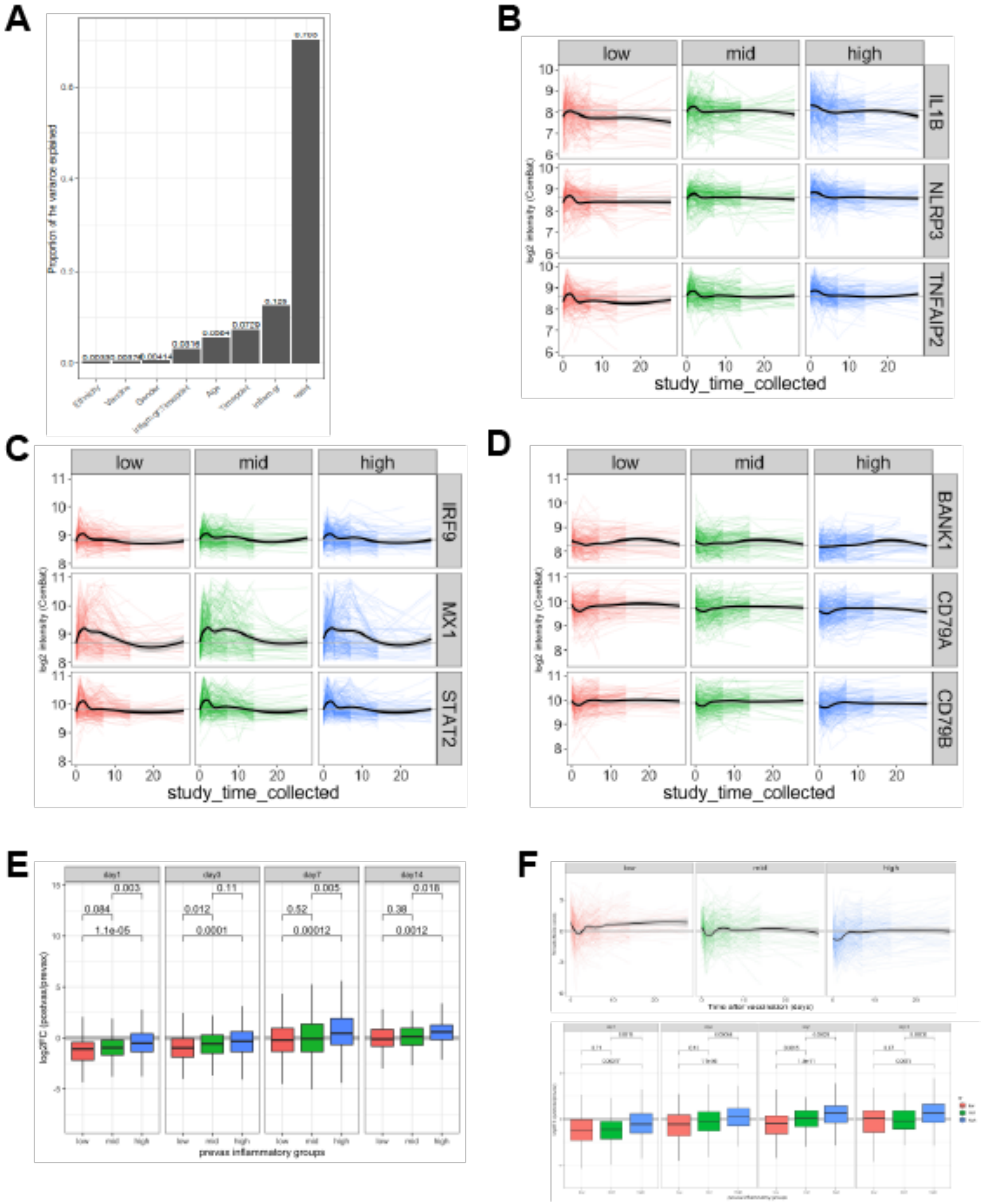


**Figure S3.** (A) Principal variance analysis with the inflammatory states. (B-D) Canonical inflammatory (B), interferon stimulated genes (C) and B cell markers (D). (E) Log2 fold-change over pre-vaccination levels of B cell markers. (F) Th2 cell markers over time (top) and fold-change over pre-vaccination (bottom).

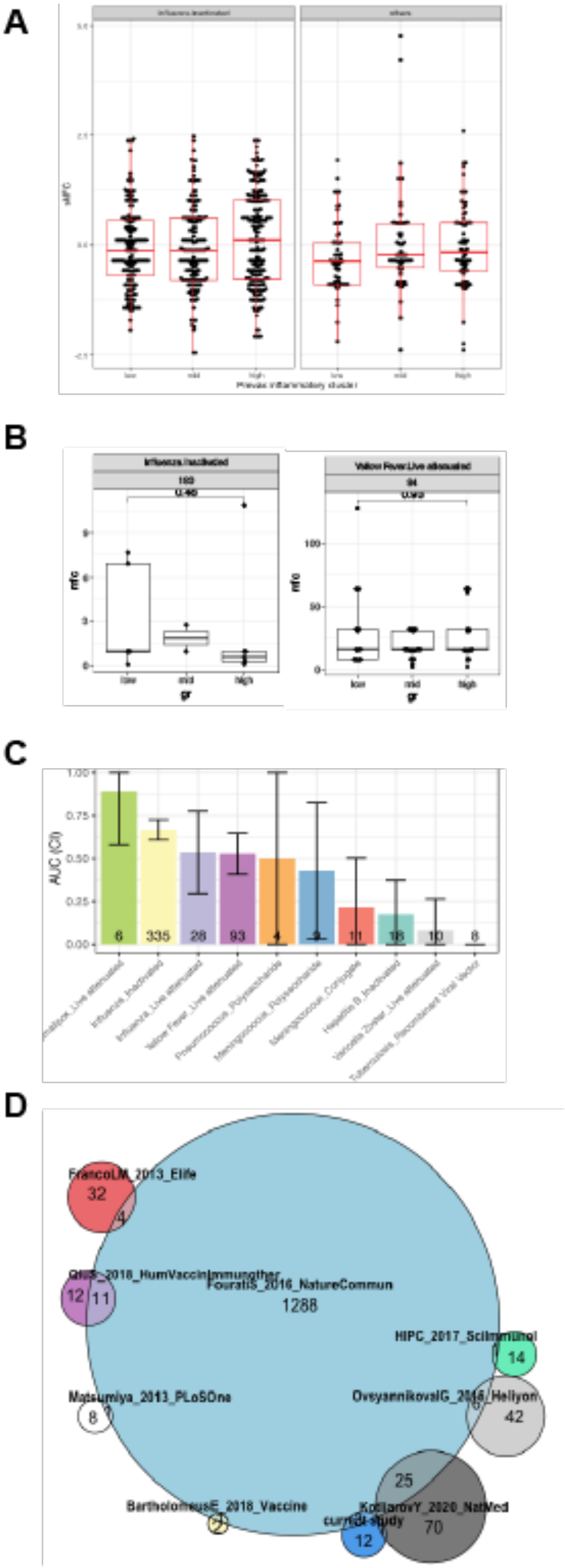
A
low
mid
high
B
Influenza.Inactivated
Yellow Fever.Live attenuated
mfc
gr
low
mid
high
C
AUC (CI)
1.00
0.75
0.50
0.25
0.00
6
335
28
93
4
9
11
18
10
8
D
FrancoLM_2013_Elife
32
4
FouratiS_2016_NatureCommun
1288
12
11
HIPC_2017_SciImmunol
14
Matsumiya_2013_PLoSOne
8
OvsyannikovaIG_2016_Heliyon
6
42
25
BartholomeusE_2018_Vaccine
2
KotliarovY_2020_NatMed
70
current study
12

**Figure S4. (A)** Association between the unsupervised pre-vaccination cluster and antibody response at day 28 for (right) Influenza inactivated vaccines (left) other vaccines. **(B)** Association between the unsupervised pre-vaccination cluster and antibody response at day 180 for the inactivated influenza vaccine (left) and day 84 yellow fever vaccine (rightt)*. **(C)*** Accuracy of the supervised classifier to predict the antibody response groups per vaccine. Mean and 95%CI for AUC across the 10-folds for each vaccine separately. **(D)** Venn diagram of the overlap of inflammatory genes and previously identified pre-vaccination signature of vaccine response.

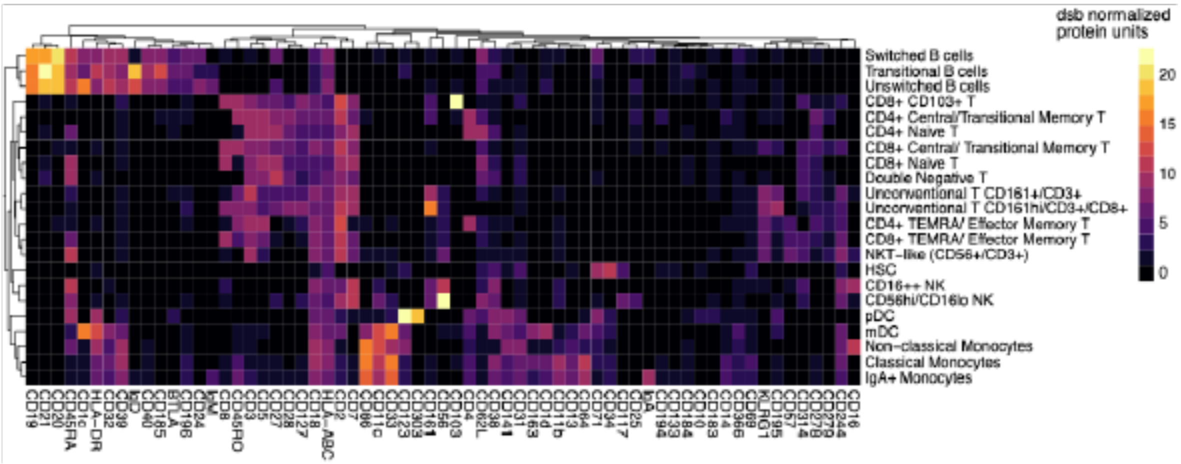


**Figure S5** PBMCs from 10 high and 10 low responders profiled by CITE-seq[1]; subsets were identified based on surface protein expression (average dsb normalized protein expression within each cluster).

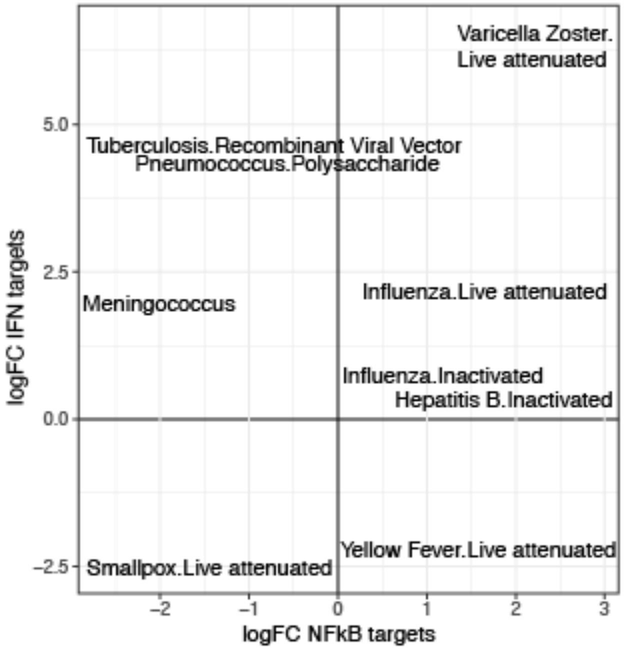

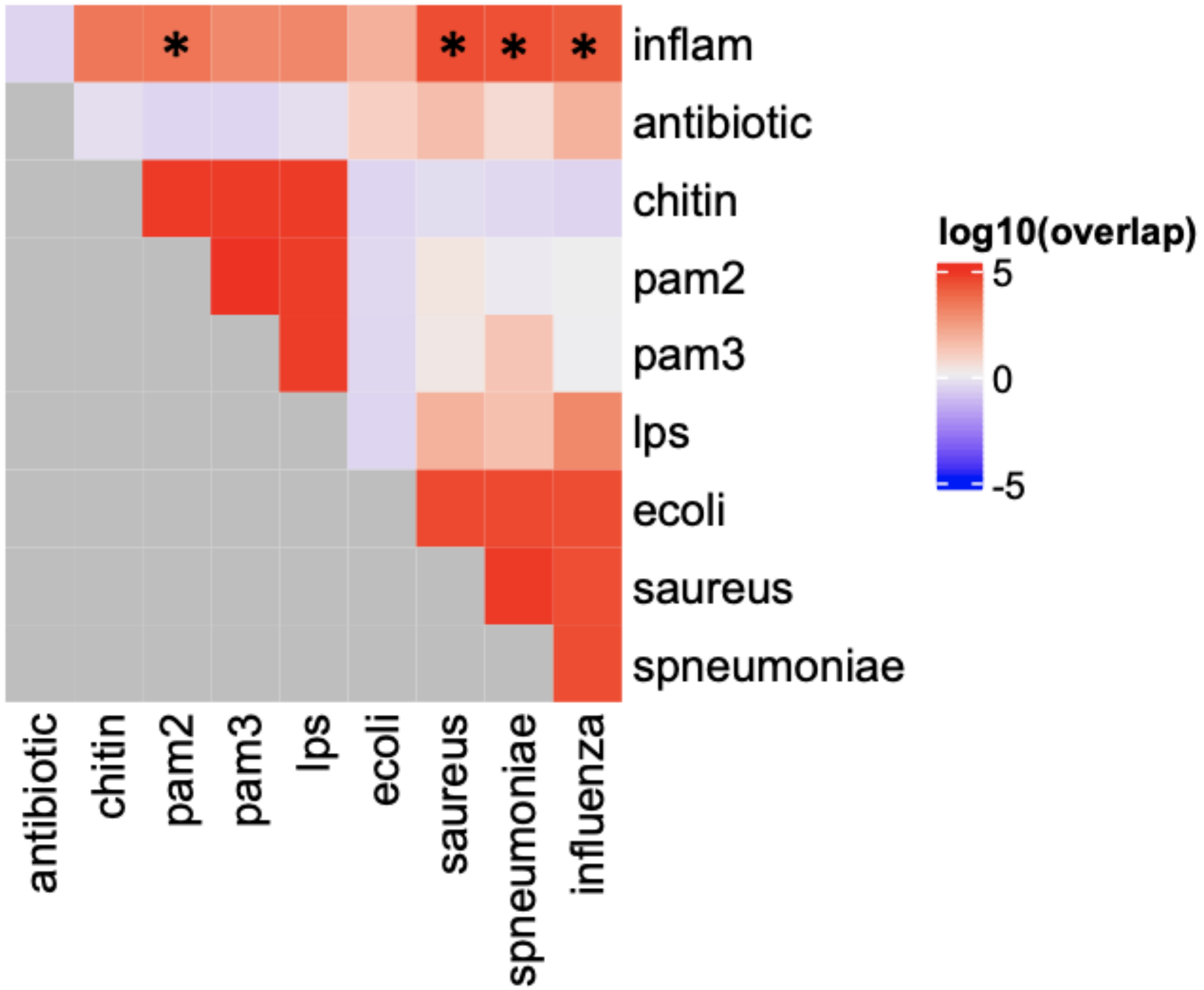


**Figure S6 (A)** Scatter plot showing the discriminative power of NFκB- and Interferon-target genes for the different vaccine present of the compendium. LogFC of the SLEA z-score of the two genesets between high vaccine responders and low vaccine responders is shown. **(B)** Overlap between the genes differentially expressed between the high and low inflammatory states and inflammatory signatures described in the literature. The significance of the overlap between ranked lists of genes was assessed by permutation.

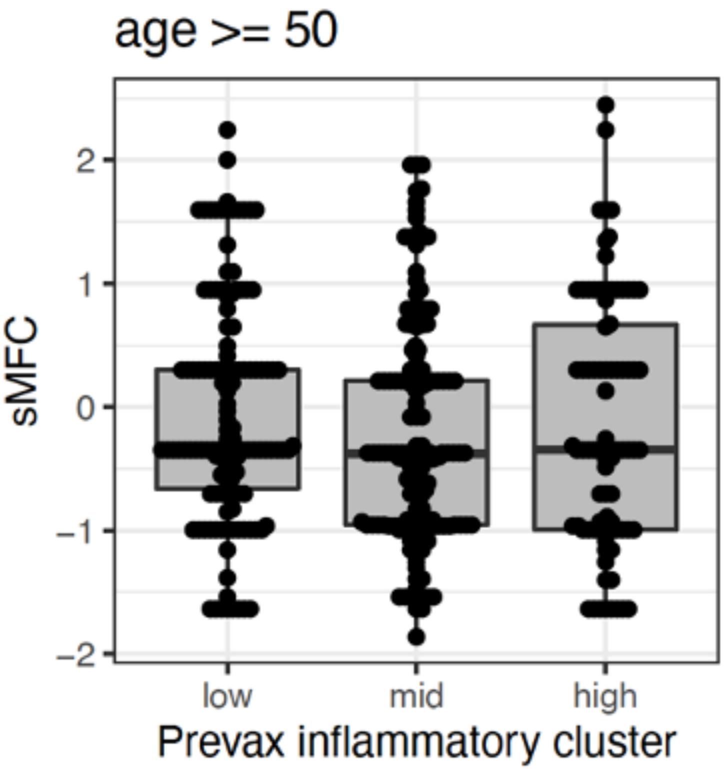


**Figure S7** Association between the unsupervised pre-vaccination cluster and antibody response at day 28 for healthy adults aged 50 and above.
